# Interactions between the N- and C- termini of mechanosensitive ion channel *At*MSL10 are consistent with a three-step mechanism for activation

**DOI:** 10.1101/726521

**Authors:** Debarati Basu, Jennette M. Shoots, Elizabeth S. Haswell

## Abstract

Although a growing number of mechanosensitive ion channels are being identified in plant systems, the molecular mechanisms by which they function are still under investigation. Overexpression of the mechanosensitive ion channel MSL (MscS-Like)10 fused to GFP triggers a number of developmental and cellular phenotypes including the induction of cell death, and this function is influenced by seven phosphorylation sites in its soluble N-terminus. Here, we show that these and other phenotypes required neither overexpression nor a tag and could be also induced by a previously identified point mutation in the soluble C-terminus (S640L). The promotion of cell death and hyperaccumulation of H_2_O_2_ in *35S:MSL10^S640L^-GFP* overexpression lines was suppressed by N-terminal phosphomimetic substitutions, and the soluble N- and C-terminal domains of MSL10 physically interacted. We propose a three-step model by which tension-induced conformational changes in the C-terminus are transmitted to the N-terminus, leading to its dephosphorylation and the induction of adaptive responses. Taken together, this work expands our understanding of the molecular mechanisms of mechanotransduction in plants.

**HIGHLIGHT:** Cell death is triggered by mutations in either the cytoplasmic N- or C-terminus of *AìMSLlü.* Our proposed model explains how membrane tension may activate signaling through the interaction of these two domains.

## INTRODUCTION

Mechanical forces, whether endogenous or exogenous in origin, are important stimuli that help direct plant growth, development, and immune responses. External mechanical perturbations include touch, wounding, wind, vibration, gravity, osmotic stress, and pathogen invasion (Monshausen and Haswell, 2013; Jayaraman et al., 2014). Endogenous mechanical forces include turgor pressure, a fundamental cue influencing cell expansion (Hamant and Haswell, 2017; Kierzkowski, 2019). Before plants can coordinate a cellular response to these varied stimuli, they must be able to precisely sense and transduce mechanical cues.

Mechanosensitive (MS) ion channels provide a fast and efficient molecular mechanism for transducing mechanical stimuli into intracellular signals. They are found in all domains of life, and are highly diverse in terms of their structure, ion selectivity, regulation, and physiological roles (Martinac et al., 2013; Ranade et al., 2015; Basu and Haswell, 2017). However, all MS ion channels share two common attributes: opening in response to mechanical force (lateral membrane tension, forces relayed from the cytoskeleton, or a combination of both) and facilitating the passive flow of ions across membranes (Bavi et al., 2017).

In plants, there is a growing list of predicted and established MS ion channel families, which includes the MscS-Like (MSL, Basu and Haswell, 2017), Two Pore Potassium (TPK, Maathuis, 2011), Mid1-Complementing Activity (MCA, Nakagawa et al., 2007) and Reduced Hyperosmolality-induced [Ca^2+^] Increase (OSCA, (Hou et al., 2014; Yuan et al., 2014; Murthy et al., 2018) families, as well as homologs of the animal MS ion channel Piezo (Zhang et al., 2019). Plant MS channels have been implicated in a wide range of physiological functions, including organellar osmoregulation (Veley et al., 2012), stomatal closure (Gobert et al., 2007; Yuan et al., 2014), cold tolerance (Mori et al., 2018), hyperosmolarity-evoked Ca^2+^ influx (Yuan et al., 2014), penetration of hard substrates (Nakagawa et al., 2007), pathogen-triggered immunity (Zhang et al., 2017) and viral resistance (Zhang et al., 2019). However, the detailed molecular mechanisms by which MS channels participate in these events need further investigation.

The MSL family of MS ion channels was first identified based on homology to the canonical MS ion channel from *Escherichia coli,* <u>M</u>echanosensitive ion <u>c</u>hannel of <u>S</u>mall conductance (*Ec*MscS) (Pivetti et al., 2003; Haswell and Meyerowitz, 2006; Haswell et al., 2008). There are ten MSLs encoded in the *Arabidopsis thaliana* genome, and they are localized to mitochondria (MSL1, Lee et al., 2016), to chloroplasts (MSL2/3, Haswell and Meyerowitz, 2006) or to the plasma membrane (MSL8, MSL9 and MSL10, Haswell et al., 2008; Hamilton et al., 2015). The tension-gated ion flux of MSL1, MSL8, and MSL10 have been examined by single channel patch-clamp electrophysiology (Maksaev and Haswell, 2012; Hamilton et al., 2015; Lee et al., 2016). MSL2/MSL3 and MSL8 are thought to serve as osmotic safety valves in plastids and in pollen, respectively (Veley et al., 2012; Hamilton et al., 2015). MSL1 plays a poorly understood role maintaining redox homeostasis in mitochondria (Lee et al., 2016).

While MSL10 was the first in the family to be characterized by electrophysiology, its physiological function is still under study. To date, no visible loss-of function phenotypes have been reported in *msl10-1* null mutants (nor in double *msl9-1 msl10-1* mutants, where MSL10’s closest homolog MSL9 is also ablated). Overexpression of *MSL10-GFP* results in growth retardation, ectopic cell death, constitutive production of H_2_O_2_, and induction of genes involved in ROS accumulation, senescence, and abiotic and biotic stress responses (Veley et al., 2014). These gain-of-function phenotypes can be attributed to a single domain of MSL10, the soluble N-terminus; in a transient *Nicotiana benthamiana* expression system, this domain can induce cell death on its own (Veley et al., 2014). In another study, we showed that variants of MSL10-GFP harboring pore-blocking lesions also induce cell death in this system (Maksaev et al., 2018). None of the phenotypes associated with *MSL10-GFP* overexpression are observed in plants overexpressing a version of *MSL10-GFP* harboring four phospho-mimetic substitutions in the N-terminus (*MSL10 ^S57D, S128D, S131E, T136D^-GFP*, or *MSL10^4D^-GFP*), though this variant has MS ion channel activity, protein levels, and subcellular localization indistinguishable from the wild type (Veley et al., 2014).

Thus, dephosphorylation of the MSL10 N-terminus activates *MSL10-GFP* to trigger cell death when overexpressed in Arabidopsis or in *N. benthamiana.* Ion flux through the channel pore, however, is not required. We infer that under normal conditions, MSL10 remains in its inactive state (i.e. with a phosphorylated N-terminus), and that overexpression overwhelms the kinase that normally maintains it in its inactive form, thereby triggering a cell death signaling cascade. It has been shown that the opening of *Ec*MscS is accompanied by structural rearrangement of the soluble C-terminal domain (Bass et al., 2002; Wang et al., 2008; Machiyama et al., 2009; Rowe et al., 2014). If similar tension-induced rearrangements of the C-terminus occur in MSL10, they might be conveyed to the N-terminus, and thereby lead to active signaling.

Indeed, a functional link between the N- and the C-termini of MSL10 was suggested by a recent report by the Zhou group (Zou et al., 2015). The *rea1 (<u>R</u>AP2.6 <u>e</u>xpresser in <u>s</u>hoot <u>a</u>pex)* mutant harbors an EMS-induced point mutation (S640L) located in the soluble C-terminus of *MSL10* that leads to increased expression of a wound-responsive luciferase reporter gene. Mutant *rea1* plants exhibit growth retardation and ectopic cell death, reminiscent of *MSL10-GFP* overexpression lines. The *rea1* mutants also exhibited shorter petioles, accumulation of anthocyanin pigments, lack of apical dominance, wound-induced hyperaccumulation of JA, and altered expression of genes involved in JA biosynthesis and response (*LIPOXYGENASE2* or *LOX2, PLANT DEFENSIN* or *PDF1.2,* and *ALLENE OXIDASE* or *AOS).*

The *rea1* mutant thus provides the opportunity to probe MSL10 function in an endogenous context. Its phenotypic similarity to *MSL10-GFP* overexpression lines suggested that growth retardation, ROS accumulation, and cell death might reflect physiological functions of the MSL10 protein, though it was also possible that these similarities might be only superficial, or that the presence of certain C-terminal tags might produce altered transgene function (Zou et al., 2015). As a result, we made an in-depth comparison of the developmental, cellular and gene expression phenotypes resulting from *MSL10-GFP* overexpression, the *rea1* lesion, and genomic phospho-variants of *MSL10.* The results from these experiments, indicating similar phenotypes among all these gain-of-function alleles, prompted us to assess genetic and physical interactions between the soluble N- and C-termini of MSL10. Our results are consistent with a three-step process of MSL10 activation that transduces the effect of increased membrane tension from the soluble C-terminus to the N-terminus of the channel, leading to phosphorylation and subsequently triggering downstream signaling and eventual cell death.

## METHODS

### Sequence alignments

The full-length amino acid sequence of *Arabidopsis thaliana* MSL10 was used as a BLASTp query to search for homologs in selected plant species. The protein with the highest identity to *At*MSL10 in each species was chosen for the final alignment, using the PRALINE multiple sequence alignment server (Bawono et al., 2017).

### Plant lines and growth conditions

The plants used in this study are all in the Arabidopsis *Col-0* ecotype background. The T-DNA insertion lines, including *msl10-1* (SALK_114626) and *msl9-1* (SALK_114361) was obtained from the Arabidopsis Biological Resource Center (Haswell et al., 2008). The *rea1/msl10-3G* mutant was obtained from the J.-M. Zhou lab (ShanghaiTech University, Shanghai). In most experiments, plants were grown on soil at 21°C under a 24-h light regime (~120 μmol m^-2^s^-1^). Backcrosses and outcrosses were made through standard techniques and genotyped with PCR-based markers. The *MSL10-GFP* overexpression lines are described in an earlier study (Veley et al., 2014).

### Genotyping

DNA was extracted by grinding tissue in extraction buffer (200 mM Tris-HCl pH 7.5, 250 mM NaCl, 250 mM EDTA, 0.5% SDS) and precipitating with an equal volume of isopropanol. The *msl10-3G/rea1* allele was genotyped by amplifying the genomic region surrounding the point mutation using gene specific oligos listed in **Supplemental Table 1** followed by digestion with *TaqI□*. restriction enzyme (NEB), which only digests the wild-type *MSL10* gene. PCR genotyping of *msl10-1* and *msl9-1* alleles was performed as described (Haswell et al., 2008).

### Cloning and generation of transgenic plants

*MSL10^S640L^* cDNA constructs were generated through site-directed mutagenesis as described (Jensen and Haswell, 2012) using pENTRY clones (pENTR+MSL10 and pENTR+MSL10^7D^) as template (Veley et al., 2014). They were cloned into pEarleyGate103 destination vectors (Earley et al., 2006) using LR recombination. To construct *gMSL10* plasmids, the *MSL10* gene was amplified from genomic wild-type *Col-0* DNA using gene-specific primers listed in **Table S1** and cloned into pENTR/D-TOPO entry vector (Thermo Fisher Scientific), then recombined into pBGW destination vectors (Karimi et al., 2002) by LR recombination. The *MSL10g^7A^* and *MSL10g^7D^* constructs were generated in a single reaction using a two-fragment Gibson Assembly, using Gibson Assembly NEB Mix and overlapping primers following manufacturer’s recommendation (Thermo Fisher Scientific). The assembled plasmids were then recombined into pBGW destination vectors by LR recombination. All primers used for Gibson cloning are listed in **Table S1**. For generating constructs for conditional expression of *MSL10* and its phospho-variants (*MSL10^4A^* and *MSL10^4D^*) under the dexamethasone (DEX)-inducible promoter, the Gateway cassette-containing region from pEarleyGate100 (Earley et al., 2006), was amplified from the plasmid introduced into the binary expression vector pTA7002 (Aoyama and Chua, 1997) using XhoI and SpeI restriction sites. pENTR constructs containing coding region of *MSL10, MSL10^4A^* and *MSL1C^4D^* (Veley et al., 2014) were used in recombination reactions with the pTA7002 vector using LR Clonase.

### Plant transformation

All binary constructs were introduced into wild-type *Col-0, msl10-1* or *msl9-1; msl10-1* plants with *Agrobacterium tumefaciens* GV3101 by floral dip (Clough and Bent, 1998). Homozygous T3 or T4 lines with a single transgene insertion were identified using selectable markers, PCR genotyping, RT-PCR, and/or fluorescent GFP expression and immunodetection.

### Gene expression analysis

Quantitative reverse-transcription polymerase chain reaction (qRT-PCR) was performed as described (Hamilton et al., 2015) with minor modifications. Total RNA was isolated from the rosette leaves of healthy 3-week-old plants (before yellow necrotic lesions develop) using the RNeasy Plant Mini Kit (Qiagen). All results shown include data from three biological replicates; for each biological replicate, three technical replicates were performed for each of three samples. Transcript levels for each gene was normalized against the geometric mean of the threshold cycle (ct) values of the two reference genes (Biazzi et al., 2015), namely *UBQ5* and *EF1α*. Finally, relative abundance of transcripts was calculated using the 2^-ΔΔ*C*t^ method (Livak and Schmittgen 2001). Semi-quantitative RT-PCR was performed as described (Veley et al., 2014). For **Figure S3A**, 10 μM DEX (Sigma-Aldrich), dissolved in 0.016% ethanol, was infiltrated into 5-week-old leaves. Tissue was harvested for RNA isolation 12 h post-infiltration. The primers used are listed in **Table S1**.

### Immunodetection

Total proteins were extracted from non-chlorotic leaves of three-week-old plants as described (Veley et al. 2014). Proteins samples were resolved by 10% SDS-PAGE and transferred to polyvinylidene difluoride membranes (Millipore) for 16 h at 100 mA. Transferred proteins were probed with anti-tubulin (Sigma, 1:10,000 dilution) and anti-GFP (Takara Bio, 1:5,000 dilution) antibodies with incubation for 1 h and 16 h, respectively. After incubation for 1 h with horseradish peroxidase-conjugated anti-mouse secondary antibodies (1:10,000 dilution; Millipore). Detection was performed using the SuperSignal West Dura Detection Kit (Thermo Fisher Scientific).

### Statistical analyses

Statistical evaluations were conducted using R Studio software (RStudio) and GraphPad Prism 8 software. Statistical differences were analyzed as indicated in the figure legends.

### Trypan blue staining and quantification

Cell death was visualized in leaves of three to four-week-old soil-grown plants using Trypan blue staining as described (Veley et al., 2014). Images were obtained with an Olympus DP80 equipped with a cooled color digital camera. The size of Trypan blue stained regions was quantified using ImageJ software as described (Fernández-Bautista et al., 2016).

### Detection and quantification of reactive oxygen species

Superoxide anion radical accumulation was detected by NitroBlue Tetrazolium chloride (NBT; Sigma) as described previously (Wilson et al., 2016). The amount of formazan was determined by measuring lysates at an absorbance of 700 nm using a 96-well microplate reader (Infinite 200 PRO; Tecan). Absorbance reads were normalized to the fresh weight of the leaves. H_2_O_2_ levels were measured by 3,3’-diaminobenzi dine (DAB, Sigma) staining (Veley et al., 2014). For the quantitative measurement of hydrogen peroxide concentrations, the Amplex Red Hydrogen/Peroxidase Assay Kit (Molecular Probes, Invitrogen) was used, following the manufacturer’s instructions and as described in (Wilson et al., 2016).

### Split-Ubiquitin Yeast two hybrid assay (mbSUS)

Intramolecular interactions between variants of the MSL10 N- and C-termini were determined using the mating-based split-ubiquitin system as described (Obrdlik et al., 2004; Lee et al., 2019). Briefly, cDNAs corresponding to the N terminal (1–460) with or without mutation of their phosphosites, the C-terminal (461-734) half of MSL10, or the soluble N- and C-terminal domains (MSL10_1-164_, MSL10_553-734_ and MSL10^S640L^_553-734_) were cloned and recombined into the destination vector pEarleyGate103 (Earley et al., 2006) and pDEST-VYCE(R)GW or pDEST-VYNE(R)GW (Gehl et al., 2009) binary vectors using LR Clonase II (Thermo Fisher Scientific). Using universal primers attB1-F and attB2-R, truncated MSL10 sequences were amplified. The primers are listed in **Table S1**. The C-terminus containing PCR products (insert) and corresponding pMetYCgate (Cub vector) were double digested with PstI+HindIII restriction enzymes, while the N-terminus containing PCR products and corresponding pXNGate21-3HA (NubG vector) were double digested with EcoRI+SmaI restriction enzymes. Subsequently, these gel-purified digested inserts and Cub or Nub vectors were co-transformed into yeast strain THY.AP4 and THY.AP5, respectively. The Cub and Nub clones were plated and selected on Synthetic Complete media lacking leucine and Synthetic Complete media lacking tryptophan and uracil, respectively. Similar to pXNGate21-3HA, N-terminus of MSL10 was also cloned into pXNGateWT (a positive control for interaction with Cub) and transformed in yeast strain THY.AP4. Diploid cells were generated by mating for two days on Synthetic Complete media lacking Leu, Trp, and Ura. After three days of growth on Synthetic Minimal media, the strength of interaction between the Nub and Cub fusions were quantified by in diploid cells using a colorimetric reporter assay with CPRG, a chlorophenol red-β-D-galactopyranoside as its substrate. All the yeast vectors were obtained from Arabidopsis Biological Resource Center. The PCR primers used for creating constructs for mbSUS assay are listed in **Table S1**. All the yeast vectors were obtained from Arabidopsis Biological Resource Center.

### Bimolecular Fluorescence Complementation (BiFC) assay

Entry vectors containing various truncated versions of the *MSL10* coding region were recombined into the binary vectors pDEST-VYCE(R)GW or pDEST-VYNE(R)GW (Gehl et al., 2009), which carry the C-terminal or N-terminal fragment of Venus YFP, respectively, using LR Clonase. The PCR primers used for cloning of BiFC constructs are listed in **Table S1**. In all cases, MSL10 fragments were tagged at the C-terminus. These plasmids were introduced into *Agrobacterium strain GV3101* and pairwise combinations were co-infiltrated into 4- to 6-week-old *N. benthamiana* leaves as described (Waadt and Kudla, 2008). To suppress posttranscriptional gene silencing, each construct pair was co-infiltrated with *Agrobacterium* strain *AGL-1* harboring p19 (Voinnet et al., 2002). Infiltrated abaxial leaf areas were examined for YFP signal using a confocal microscope (Olympus Fluoview FV 3000) at 3 to 5 d post-inoculation. The experiments were performed at least three times using different batches of plants; for each biological replicate, three independent *N. benthamiana* plants were infiltrated.

### Accession numbers

GenBank accession numbers used in **Figure 1** are: *Arabidopsis thaliana* (NP_196769.1), *Arabidopsis lyrata* (XP_002873549.1), *Brassica rapa* (XP_009121883.1), *Brassica napus* (XP_013676093.1), *Camelina sativa* (XP_010453270.1), *Populus trichocarpa* (POPTR_0006s14640g), *Medicago truncatula* (XP_003603202.2), *Vitis vinefera* (XP_002279755.1), *Solanum lycopersicum* (XP_004245056.1), *Solanum tuberosum* (XP_006350354.1), Zea *mays* (XP_008649202.1), *Oryza sativa* (XP_015641284.1), *Sorghum bicolor* (XP_002438025.1), *Brachypodium distachyon* (XP_003560953.1), *Setaria italica* (XP_004964936.1).

**Figure 1.**
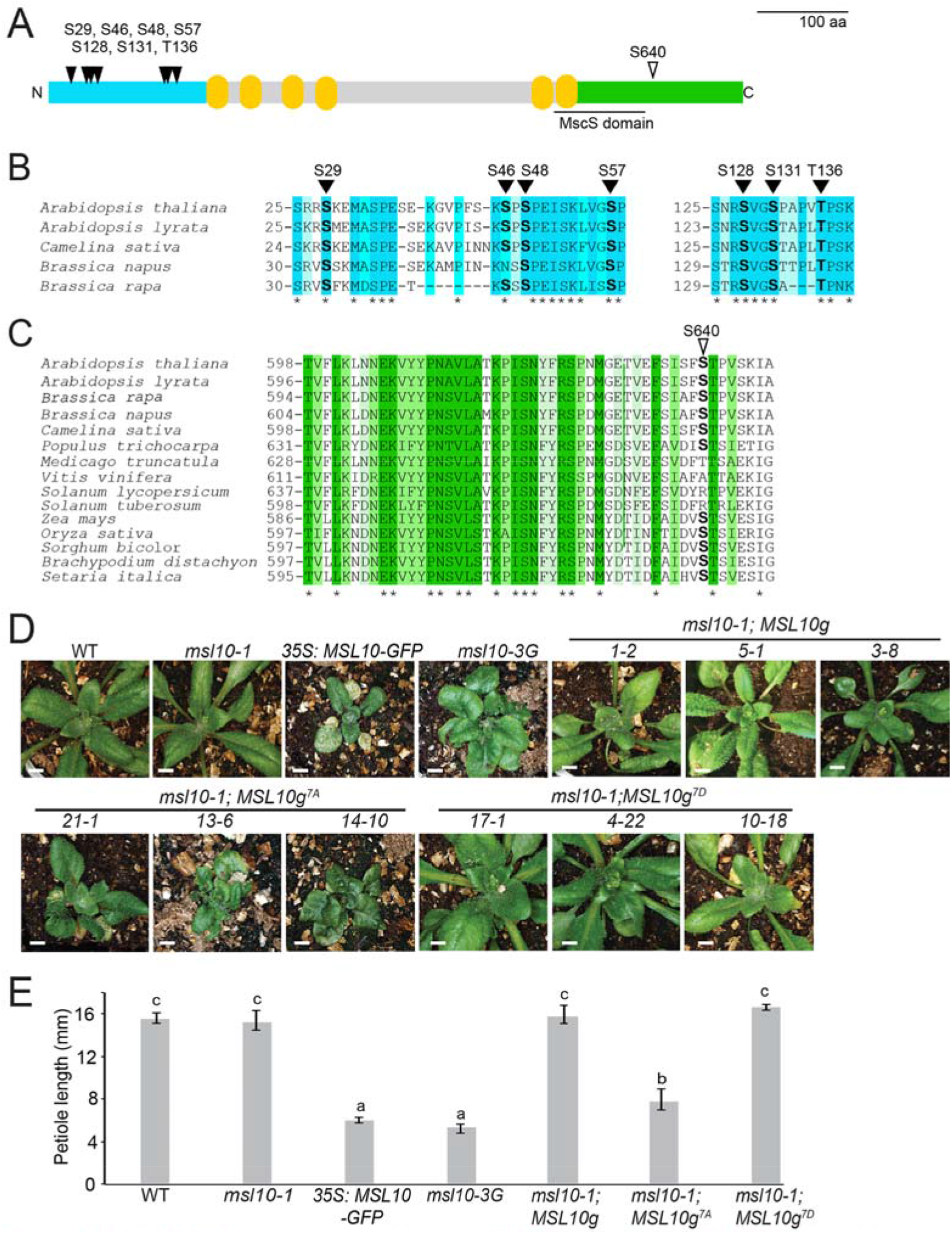
*MSL10-GFP* overexpression lines, *msl10-3G* mutants, and *MSL10g^7A^* lines have similar whole-plant phenotypes. **(A)** Predicted topology of an *At*MSL10 monomer. Teal, cytosolic N-terminus; yellow, transmembrane helices; green, cytosolic C-terminus. Black triangles denote the positions of the residues mutated in this study. The MscS domain is as defined in (Veley et al. 2014). **(B)** Multiple alignment of protein sequences corresponding to the N-terminus of putative *At*MSL10 orthologs in the *Brassicaceae* family. Darker teal shading indicates higher conservation score and identical amino acids with an asterisk. (C) Multiple alignment of protein sequences corresponding to the C-terminus of putative *At*MSL10 orthologs in selected flowering plants. Conservation of residues is indicated as in **(B). (D)** and **(E)** T4 homozygous lines overexpressing *MSL10-GFP* in the *Col-0* background (line *15-2*), T3 homozygous lines expressing *MSL10g,* or *MSL10g^7D^* in the *msl10-1* background, and T3 segregating or T3 homozygous lines expressing *MSL10g^7A^* in the *msl10-1* background were selected for comparison. **(D)** Whole-plant images of 25-day-old plants of the indicated genotypes grown side-by-side at 21°C under a 24-h light regime. Bar = 0.5 cm. **(E)** Petiole length of the fourth or fifth leaf from 28-day-old plants. Different letters indicate significant differences as determined by one-way ANOVA followed by Tukey’s post-hoc test (P < 0.05). Error bars indicate ± SD of three replicates (n = 18 per replicate).

The rest of the genes referred to in this study correspond to the following Arabidopsis Genome Initiative locus identifiers: *MSL9 (At5G19520), MSL10 (At5G12080), RAP2.6 (At1g43160), AOS (At5g42650), LOX2 (At3g45140), PDF1.2 (At5G44420), SAG12 (At3g20770), PERX34 (At3g49120), DOX1 (At3g01420), OSM34 (At4G11650), UBQ5 (At3g62250), EF1α (At5g60390).*

## RESULTS

### Conservation of functionally significant residues in the soluble N- and C-terminal domains of putative MSL10 orthologs

We investigated the conservation of eight residues previously identified as important for MSL10 function: seven phosphorylation sites in the soluble N-terminus (S29, S46, S48, S57, S128, S131, and T136; and a C-terminal amino acid that is mutated in the *rea1* mutant (S640, Zou et al., 2015) (**Figure 1A**). The predicted N-terminal domains of putative MSL10 homologs from other flowering plants showed surprisingly low sequence conservation. However, when analysis was restricted to the *Brassicacae* family, the 33-amino-acid region containing S29, S46, S48 and S57, and the 15-amino-acid region containing S128, S131, and T136 showed 73% amino acid identity with the *Brassica napus* homolog (**Figure 1B**). The sequence of the MSL10 C-terminus surrounding S640 was highly conserved among angiosperms (90% amino acid identity with the putative homolog from *B. napus,* 60% identity with that from *Oryza sativa),* and S640 itself was conserved in 11 of the 15 sequences analyzed (**Figure 1C**). To summarize, phosphosites in the soluble N-terminus known to modulate the *MSL10-GFP* overexpression phenotype were not well-conserved, while S640, a residue in the soluble C-terminus known to affect MSL10 function, was well-conserved among the sequences analyzed.

### *real* is a recessive gain-of-function *MSL10* allele, renamed *msl10-3G*

To further characterize the *rea1* allele, we backcrossed *rea1* plants to wild type *Col-0* plants. As shown in **Figure S1A**, the resulting F1 hybrid plants resembled the wild type parent, indicating that the mutation responsible for the phenotypes observed in the *rea1* mutant is recessive, consistent with previous findings (Zou et al., 2015). After self-pollination, the resulting F2 population segregated into wild-type and *rea1* phenotypes at a ratio of 3:1, as expected for a recessive allele, and these results were confirmed by PCR genotyping (**Figure S1C**). In addition, *rea1* plants were outcrossed to *msl10-1* null mutant plants. The F1 progenies resembled *rea1* in terms of leaf morphology but attained intermediate height after 8 weeks of growth (**Figure S1B**). After self-pollination, resulting F2 populations similarly segregated approximately 1:2:1 with respect to height at late stages of development (wild type like: intermediate: *rea1* phenotype); phenotypes were confirmed by PCR genotyping (**Figure S1C**). Thus, the phenotypic effects of the *rea1* lesion depend on the presence or absence of the WT *MSL10* allele, suggest that they may be dosage-dependent (as it requires two copies of the *real1* mutant allele to generate the full *rea1* phenotype), and confirm that the phenotypes associated with the *rea1* mutant are due to a recessive gain-of-function point mutation in the *MSL10* gene. This allele is hereafter called *msl10-3G*.

### *MSL10-GFP* overexpression, the *msl10-3G* allele, and the *MSL10g^7A^* transgene produce similar phenotypes and gene expression patterns

Strong constitutive overexpression of a gene often results in pleiotropic phenotypes that may not reflect its normal function (Zhang et al., 2003; Kovalchuk et al., 2013), and the addition of tags can alter protein localization, regulation, stability or function (Spartz et al., 2012). Zou et al. (2015) reported that neither estradiol-induced expression of *MSL10-FLAG* nor constitutive expression of untagged *MSL10* resulted in the phenotypes produced by constitutive expression of *MSL10-GFP,* suggesting that large tags may perturb the function of MSL10. Furthermore, we were previously unable to retrieve plants constitutively expressing *MSL10^4A^-GFP* (Veley et al., 2014), and while we proposed that the over-expression of phospho-dead *MSL10* was lethal, it remained possible that an unrelated defect in the transgene prevented us from isolating any transgenic plants.

To determine if the phenotypes associated with *MSL10-GFP* overexpression could be replicated at endogenous expression levels and without a GFP tag, we generated constructs to drive expression of untagged wild-type *MSL10 (MSL10g)*, phospho-dead *MSL10* S29A, S46A, S48A, S57A, S128A, S131A, and T136A (*MSL10g^7A^*) or phospho-mimetic *MSL10* S29D, S46D, S48D, S57D, S128D, S131E, and T136D (*MSL10g^7D^*), within the native *MSL10* genomic context. These constructs were introduced into the *msl10-1* null mutant background and three independent homozygous transgenic lines from each transformation were selected for further analysis. These lines accumulated *MSL10* transcripts at levels similar or slightly higher than those of wild-type plants (**Figure S2A**).

Like the *MSL10-GFP* overexpression lines, the *msl10-3G* mutant and three *MSL10g^7A^* lines all exhibited reduced rosette size, fresh weight and plant height compared to wild-type (**Figure 1D, S2B-D**). Four- to five-week-old plants from these same lines also exhibited shorter petioles and broader leaf blades compared to wild-type plants (**Figure 1E, S2E-F**). Unlike *MSL10-GFP* overexpression lines, the *msl10-3G* mutant and *MSL10g^7A^* lines lacked apical dominance (**Figure S2G**). The *msl10-1* null mutant and *msl10-1* mutants expressing *MSL10g* or *MSL10g^7D^* were phenotypically indistinguishable from wild-type plants.

We next tested if *msl10-3G* mutants, *MSL10^7A^g* lines, and *MSL10-GFP* overexpression lines have similar gene expression patterns. As shown in **Figure 2A**, four genes previously shown to be upregulated in *MSL10-GFP* overexpression lines (*SENESCENCE ASSOCIATED GENE 12 (SAG12), α-DIOXYGENASE (DOX1), PEROXIDASE-34 (PERX34),* and *OSMOTIN-LIKE PROTEIN-34 (OSM34)* (Veley et al., 2014) were also expressed at higher levels in *msl10-3G*(4- to 5-fold increase) and *MSL10g^7A^* lines (3- to 5-fold increase) compared to wild-type plants, although not to the same degree as in *MSL10-GFP* overexpression lines (7- to 20-fold increase). Similarly, four genes previously shown to be upregulated in *rea1* mutants, *LIPOXYGENASE2 (LOX2,) PLANT DEFENSIN (PDF1.2), ALLENE OXIDASE (AOS)* and *RAP2.6* (Zou et al., 2015) were also induced in *MSL10-GFP* overexpression lines (14- to 35-fold), and in *MSL10g^7A^* lines (5- to 12-fold) compared to the wild type (**Figure 2B**). Mutant *msl10-1, MSL10g,* and *MSL10g^7D^* lines did not exhibit statistically significant differences in expression of any of these genes compared to wild-type plants. We also observed ectopic cell death and a similar induction of *SAG12, DOX1, OSM34* and *PERX34* in response to inducible expression of wild-type and phospho-dead (but not phospho-mimetic) MSL10 (**Figure S3**).

**Figure 2.**
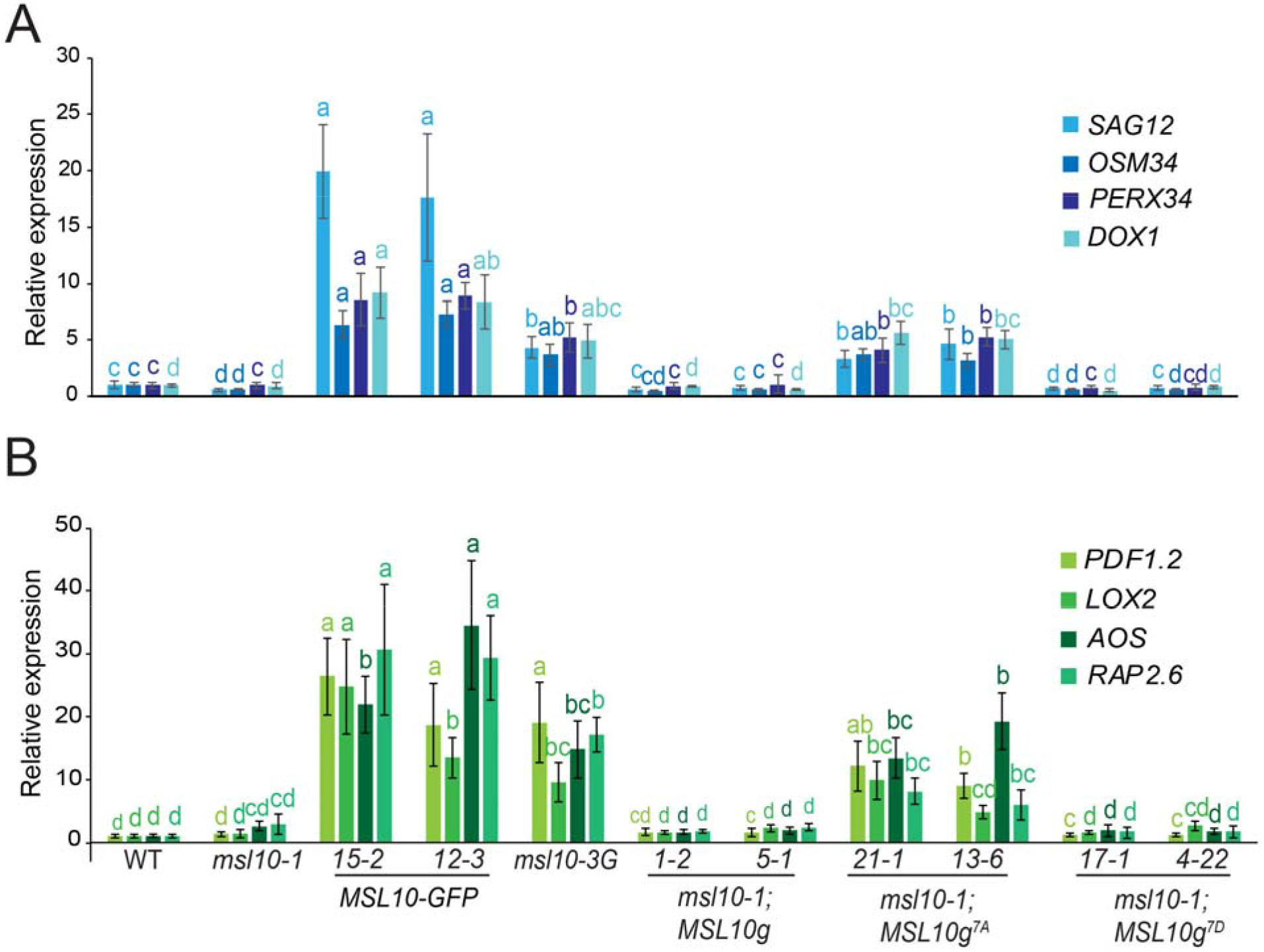
*MSL10-GFP* overexpression lines, *msl10-3G* mutants, and *MSL10g^7A^* lines display similar expression profiles. Quantitative RT-PCR analysis of four genes previously shown to be upregulated in *35S:MSL10-GFP* overexpression lines (A) and four genes previously shown to be upregulated in the *msl10-3G* background (B). cDNA was synthesized from total RNA extracted from rosette leaves of three-week-old plants grown at 21°C under 24 hours of light. Expression levels of respective genes were normalized to both *EF1a* and *UBQ5*. Mean fold-change values relative to the wild type are plotted, with error bars showing ± SE of the mean of three biological replicates Different letters indicate significant difference as determined by one-way ANOVA followed by Tukey’s post-hoc test (P < 0.05). For transgenics, two independent T3 or T4 homozygous lines were selected for comparison.

Besides these distinct morphological defects, *MSL10-GFP* overexpression lines also exhibited constitutively elevated levels of ROS and ectopic cell death (Veley et al. 2014). This prompted us to investigate whether similar hyper-accumulation of ROS and cell death were displayed by the *MSL10g^7A^* or *msl10-3G* lines. To examine superoxide radical (O^2-^) and hydrogen peroxide (H_2_O_2_) content, we employed nitroblue tetrazolium (NBT) and 3,3’-deaminobenzidine (DAB) staining of three-week old rosette leaves, respectively (Myouga et al. 2008, Wilson et al. 2016). An Amplex Red-coupled fluorescence assay was performed for quantifying H_2_O_2_ levels. *MSL10-GFP* overexpression lines, the *msl10-3G* allele, and lines expressing *MSL10g^7A^* all displayed hyperaccumulation of both superoxide and H_2_O_2_, but not *msl10-1, MSL10g* and *MSL10^7D^* lines (**Figure S4**). *MSL10-GFP* overexpression lines, the *msl10-3G* allele, and lines expressing *MSL10g^7A^* all displayed ectopic cell death, as assessed by Trypan blue staining, compared to wild-type plants. Ectopic cell death was not observed in *msl10-1, MSL10g* and *MSL10^7D^* lines (**Figure S5**).

Thus, overexpression of *MSL10-GFP,* expression of untagged *MSL10^7A^* at endogenous levels in the *msl10-1* background, and the C-terminal point mutation in the *msl10-3G* allele all lead to growth retardation, defects in petiole length, overaccumulation of ROS, and the upregulation of eight hallmark genes. While the *MSL10-GFP* overexpression line *15-2* was the most severely affected in all cases, and the only line to display yellowish-brown patches on rosette leaves (**Figure 1D**), all of these phenotypes could be generated to various degrees of severity without overexpression and/or a C-terminal tag, and thus are likely to be related to the normal function of MSL10. Furthermore, these phenotypes are a direct, rather than developmental, effect of ectopic MSL10 activation.

We interpret these data to mean that high levels of wild-type MSL10, basal levels of MSL10 that is dephosphorylated at the N-terminus, or basal levels of MSL10 harboring the S640L lesion all lead to a set of pleiotropic phenotypes through the same, as yet unknown, molecular mechanism. A lack of MSL10-specific antibodies prevented us from measuring protein levels in untagged lines, so it is formally possible that MSL10 phospho-dead lesions or the S640L lesion alter protein stability in the absence of a GFP tag. However, these lesions do not alter stability when fused to a GFP tag (see Veley et al., 2014 and **Figure 3B** below).

**Figure 3.**
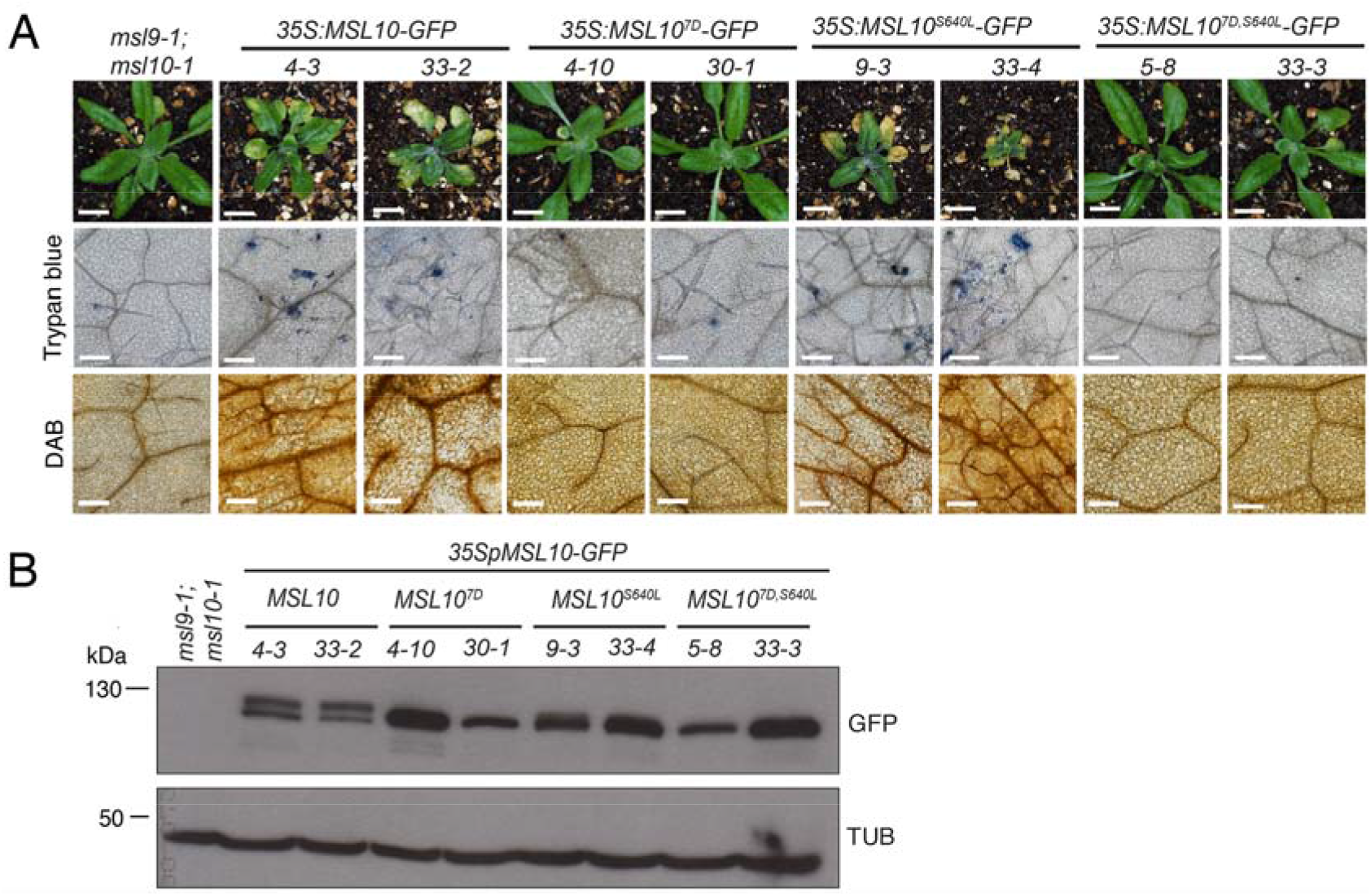
Phospho-mimetic substitutions in the MSL10 N-terminus suppress phenotypes associated with overexpression of MSL10^S640L^. **(A) Top row**: Images of three-week-old plants grown at 21°C overexpressing wild-type *MSL10-GFP*, phospho-mimetic *MSL10^7D^-GFP, MSL10^S640L^-GFP*, or *MSL10^7D,S640L^-GFP*. Overexpression lines were generated in the *msl9-1;msl10-1* background, phenotypically indistinguishable from *msl10-1*. Two independent homozygous T3 lines for each transgene are shown. Bar = 0.5 cm. **Middle row**: Trypan blue staining of four-week old leaves from the above T3 plants to visualize ectopic cell death. **Bottom row**: DAB staining of five-week-old leaves to detect the accumulation of H_2_O_2_. For leaf images, bar = 0.2 mm. **(B)** Immunoblot analysis of MSL10-GFP variants in rosette leaves of two-week old plants. MSL10-GFP was detected with an anti-GFP primary antibody (top), and then the blot was stripped and re-probed with an anti-a-tubulin primary antibody (bottom). Expected protein sizes are indicated at the left according to a commercially available standard. The two forms of MSL10-GFP that migrate slower on SDS-PAGE may result from posttranslational modifications.

### Phospho-mimetic lesions in the MSL10 N-terminus suppress *msl10-3G* phenotypes

Given that mutations in the N-terminus (*MSL10g^7A^)* and the C-terminus (*msl10-3G)* produce similar phenotypes, we tested for a genetic interaction between these two soluble domains. The *msl10-3G* (S640L) mutation was introduced into the *35S:MSL10-GFP* and *35S:MSL10^7D^-GFP* transgenes to make *MSL10^S640L^-GFP* and *MSL10^7D,S640L^-GFP*. As expected, overexpression of *MSL10-GFP* and *MSL10^S640L^-GFP* lead to growth retardation, ectopic cell death (as assessed by the occurrence of yellowish-brown lesions on rosette leaves and Trypan blue staining), and enhanced H_2_O_2_ accumulation in rosette leaves, while overexpression of phospho-mimetic *MSL10^7D^-GFP* did not (**Figure 3A**). *MSL10^7D,S640L^-GFP* plants were indistinguishable from wild type or *MSL10^7D^-GFP* plants. Immunoblotting showed that these phenotypic differences cannot be attributed to differences in protein abundance (**Figure 3B**), and likely reflect the inability of MSL10^7D^-GFP and MSL10^7D,S640L^-GFP to activate downstream signaling. Thus, N-terminal phospho-mimetic substitutions prevent or block the phenotypes produced by the C-terminal lesion S640L found in the *msl10-3G* mutant.

### The soluble N- and C-termini of MSL10 interact directly in two protein-protein interaction assays

To assess whether this genetic interaction between the N- and C-termini of MSL10 might be mediated through physical interactions, we first employed the mating-based split-ubiquitin yeast two-hybrid assay (mbSUS) (Reinders et al., 2002). We previously showed that MSL10 can interact with itself in this assay, but not with close relative MSL9—as expected for a homomeric channel (Veley et al., 2014). We repeated these results and further observed that MSL10 does not interact with an unrelated membrane protein, KAT1 (Obrdlik et al., 2004) (**Figure 4A**).

**Figure 4.**
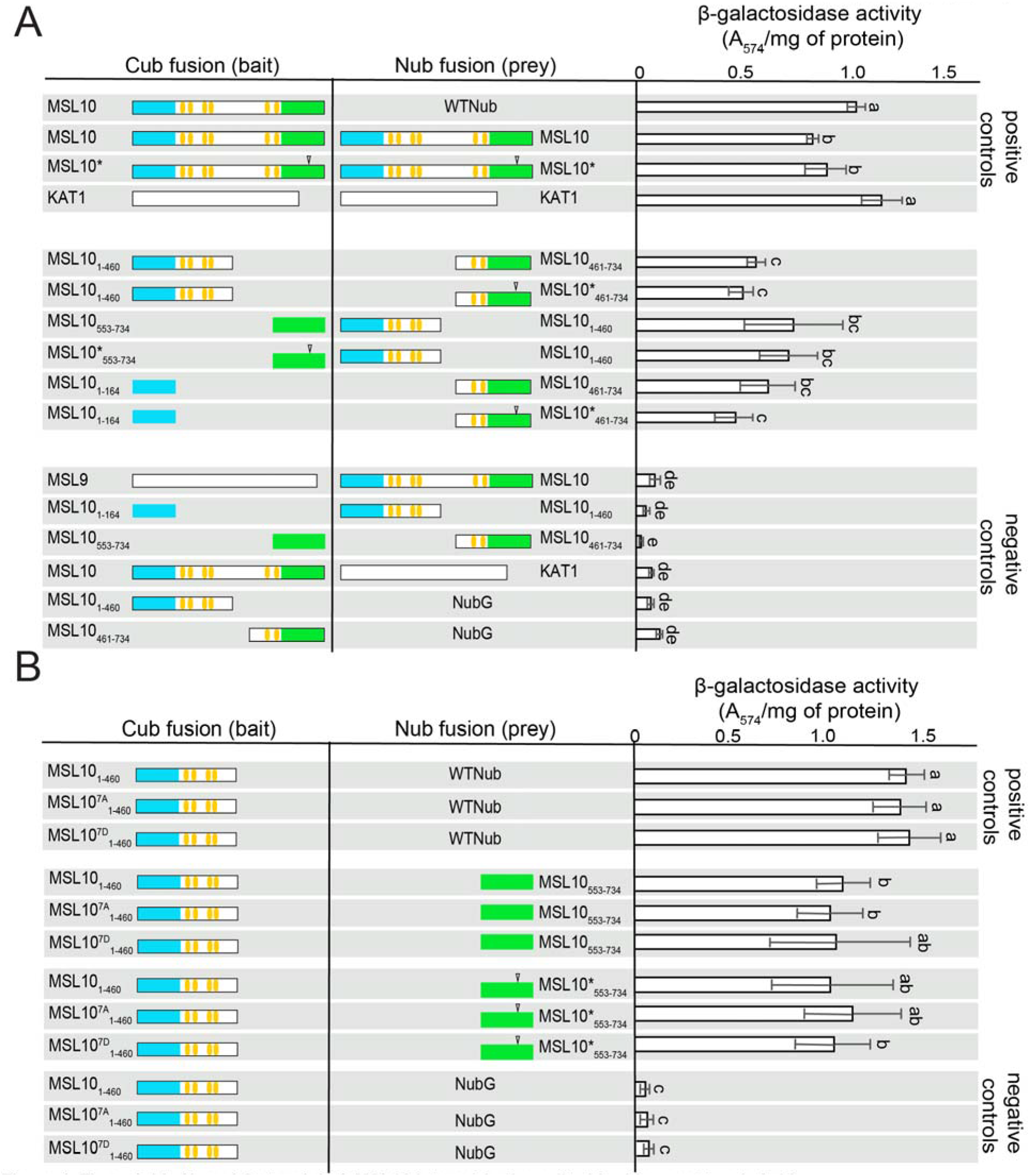
The soluble N- and C-termini of MSL10 interact in the split ubiquitin yeast two hybrid. **(A)** Specific interactions between the N- and C-termini of MSL10 do not require TM helices and are unaffected by the S640L lesion. **(B)** Phospho-mimic or phospho-dead lesions in the N-terminus do not affect interactions with the soluble C-terminus. Left and middle panels indicate fusions with the C- and N-terminal domains of ubiquitin (Cub and Nub, respectively). Teal, cytosolic N-terminus; yellow, transmembrane helices; green, cytosolic C-terminus. Asterisks and open arrows indicate the S640L lesion. Right panel, results from liquid assay for B-galactosidase activity. Data presented are means ± SD of three replicates. Different letters indicate significant differences as determined by one-way ANOVA followed by Tukey’s post-hoc test (P < 0.05).

To test for interactions specific to the N- and C-termini, we first split MSL10 into two halves, the first comprising the N-terminal domain and the first four TM helices (MSL10_1-460_), and the second comprising the fifth and sixth TM helix as well as the C-terminal domain (MSL10_461-734_). These two halves of MSL10 displayed a strong interaction, providing support for intra- or inter-molecular interactions between different domains of MSL10 (**Figure 4A**). Furthermore, the soluble N-terminus (MSL101-164) and the C-terminal half of MSL10 (MSL_10461-734_) interacted, as did the N-terminal half of MSL10 (MSL10_1-460_) and its soluble C-terminus (MSL_10553-734_). This result showed that the middle part of the protein—which contains all the TM helices—is not required for self-association. Almost no interaction was detected between the N-terminal half and itself or between the C-terminal half and itself. None of these interactions were appreciably affected by the presence of the S640L lesion (indicated as MSL10*, **Figure 4A**), phospho-mimic or phospho-dead residues (**Figure 4B**), nor by any combination of these N- and C-terminal lesions (**Figure 4B**).

To validate these mbSUS interactions, to investigate whether they can occur *in planta,* and to assess interactions without requiring one partner to be tethered to the membrane, we employed a bimolecular fluorescence complementation (BiFC) assay as described in Gehl et al. (2009). The N- and C-terminal halves of YFP (YN- and YC-) were fused to MSL10 variants and transiently expressed in leaf epidermal cells of *Nicotiana benthamiana*. As expected, co-infiltration of MSL10-YN with MSL10-YC resulted in strong YFP signal at the periphery of the cell (**Figure 5**). Strong YFP fluorescence was also detected when soluble MSL10_553-734_-YN was co-infiltrated with soluble MSL10_1-164_-YC or with the N-terminal half of the protein, MSL10_1-460_-YC (**Figure S6A**). The C-terminal half of MSL10 formed aggregates (Figure S6B) so we only used the soluble C-terminus (MSL10_553-734_) in this experiment. Neither the N-terminus nor the C-terminus of MSL10 was able to self-associate in this assay, as only patchy, diffuse signal-- similar to that observed with unfused YN or YC--was observed in these cases. Furthermore, neither of these domains were able to interact with MSL9 or KAT1 (**Figure 5, Figure S6A**). Consistent with the mbSUS assay results, the S640L mutation (again indicated as MSL10*) did not affect any interactions; nor did the introduction of phospho-mimetic or phospho-dead lesions into the N-terminal sequences. These results demonstrate a direct physical interaction between the soluble N- and C-termini of MSL10 and show that it does not require tethering to the membrane and is unaffected by the S640L lesion or by the phosphorylation status of the N-terminus.

**Figure 5.**
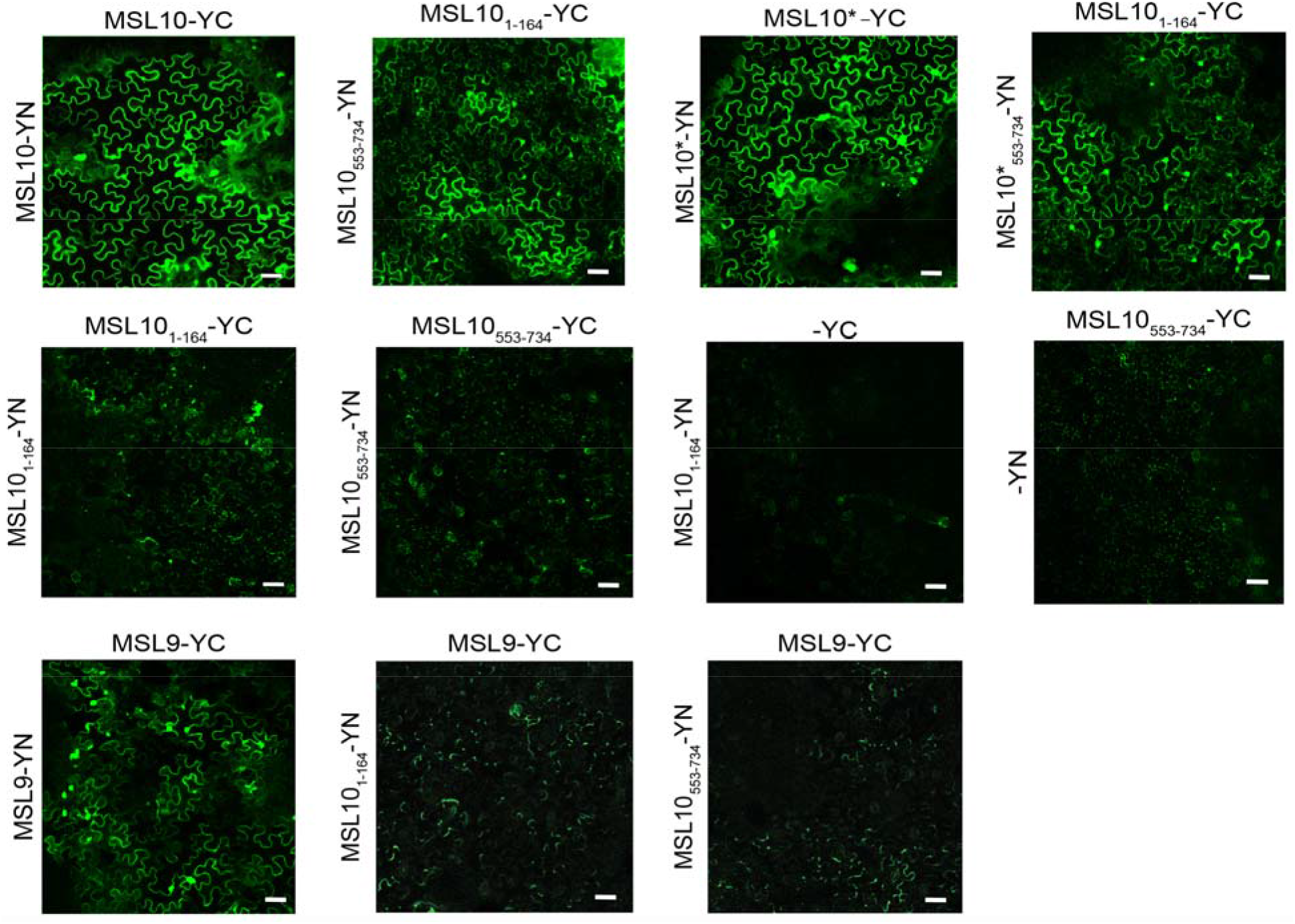
The soluble N- and C-termini of MSL10 interact in the bi-molecular fluorescence complementation (BiFC) assay. Confocal laser scanning micrographs of the abaxial surface of *N. benthamiana* leaves 3 to 5 days after infiltration with *Agrobacterium* harboring the indicated BiFC construct pairs. Scale bar, 50 μm. Identical acquisition settings were used in all images. All experiments were repeated at least three times. MSL10* indicates MSL10^S640L^.

## DISCUSSION

Here we present several lines of evidence in support of a direct interaction between the soluble N-terminus and the soluble C-terminus of the plant mechanosensitive ion channel *At*MSL10. First, a suite of phenotypes was produced by overexpression of *MSL10-GFP*, by native expression of phospho-dead substitutions in MSL10’s N-terminus, or by the S640L point mutation in its C-terminus (**Figure 1, 2, S2, S3, S4 and S5**). Second, phospho-mimetic substitutions suppressed the effect of S640L when both types of lesions were present in the same monomer (**Figure 3**). Third, the soluble N- and C-termini of MSL10 interacted in two protein- protein interaction assays in a manner independent of phospho-variant mutations or the S640L lesion (**Figure 4 and Figure 5**).

To date, functional characterization of MSL10 has been based on overexpression of *MSL10-GFP* in *N. benthamiana* and Arabidopsis (Veley et al., 2014). Here, we show that two additional types of *MSL10* gain-of-function lines (plants expressing endogenous levels of untagged *MSL10g^7A^* or harboring the *msl10-3G* allele) phenocopy *MSL10-GFP* overexpression lines, as does inducible overexpression of MSL10^7A^ (**Figure 1, 2, S2, S3**). Thus, the ability of MSL10 to induce growth retardation, ectopic cell death, hyperaccumulation of H_2_O_2_, and the induction of genes related to ROS, and cell death does not require overexpression or the presence of a tag, and therefore is likely to be related to the normal function of MSL10. We note that plants overexpressing untagged wild type *MSL10* do not show these phenotypes (Zou et al., 2015). Perhaps, in this context, a large tag improves MSL10 stability.

We thus hypothesize that the ability to trigger cell death is one of the functions of MSL10. But since ectopic cell death is only seen when endogenously expressed MSL10 is mutated, it could be claimed that the cell death we observe does not reflect of the normal function of MSL10, but rather is a non-specific toxicity associated with overexpression of mutants. The results presented here with the *msl10-3G* mutation (*MSL10^S640L^*) show that this is unlikely. First, the MSL10^S640L^ mutation does not trigger cell death when the 7D substitutions are introduced into the same monomer (**Figure 3A**), which argues against the MSL10^S640L^ mutation being inherently toxic. Secondly, the *msl10-3G* allele is recessive (Zou et al., 2015, and **Figure S1A**), and therefore is not exerting a general toxic effect. Finally, it is difficult to imagine how lesions in the N-terminus, the C-terminus, and overexpression would all have the same non-specific and toxic effect. Our current studies investigate whether increased membrane tension can cause WT MSL10 to trigger cell death, as hypothesized in **Figure 6A**, to more firmly establish cell death as a function of MSL10.

**Figure 6.**
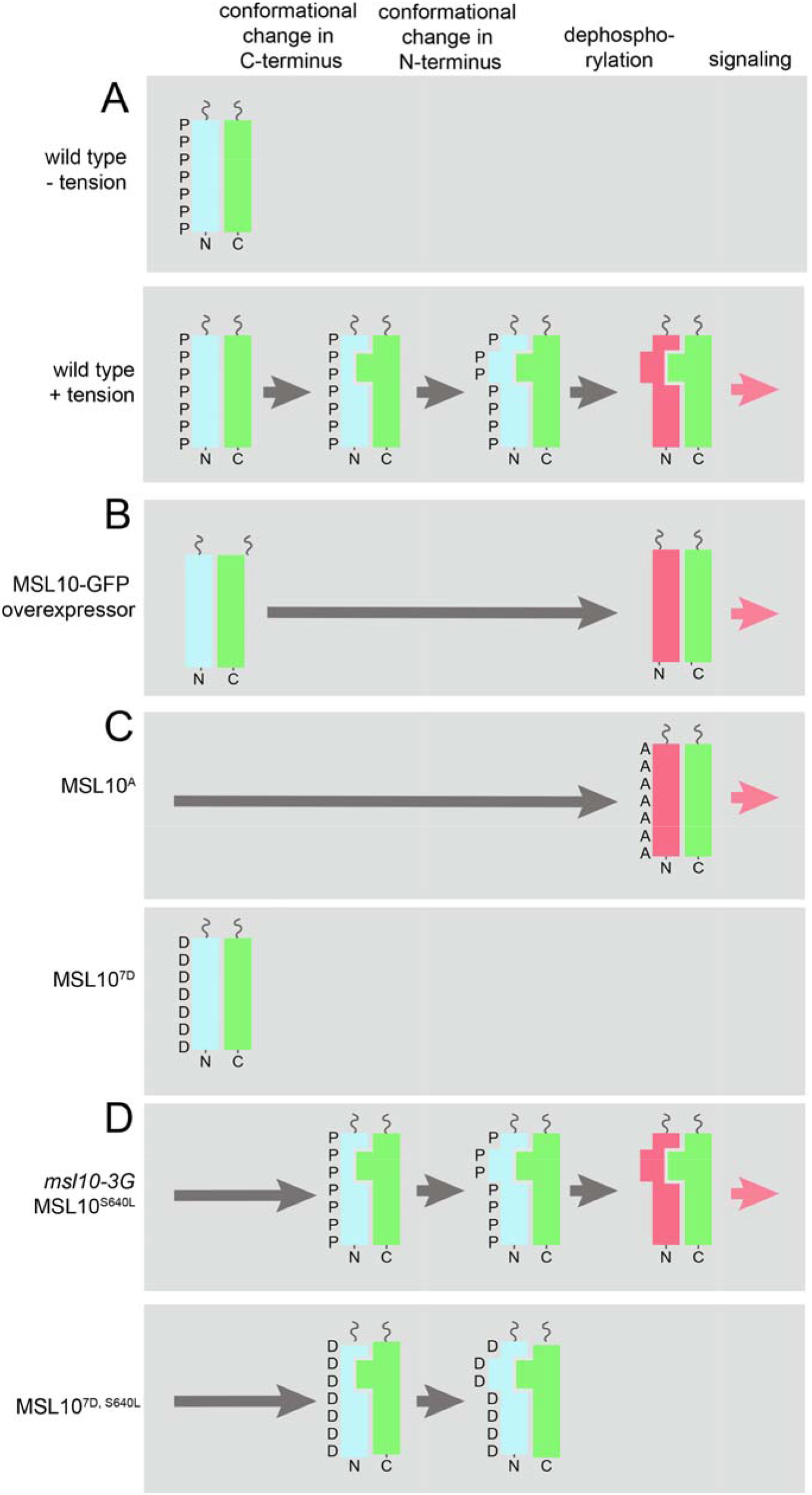
A three-step model for the regulation of MSL10’s cell death signaling function. For simplicity, only the soluble N- and C-terminal domains of MSL10 are shown. They could be from the same or from different monomers. The inactive N-terminus, active N-terminus, and C- terminus are represented by rectangles colored blue, red, and green, respectively. Proposed conformational changes are indicated by changes in the shape of each rectangle. This model explains the signaling output (red arrow) observed in wild type plants **(A)**, in response to MSL10-GFP overexpression **(B)**, in the presence of MSL10 phospho-variants **(C)**, in the presence of alleles or transgenes harboring the S640L mutation in the C-terminus (**D**, top), or in the presence of a transgene that combines S640L with phospho-mimetic substitutions in the N- terminus (**D**, bottom). See main text for more detailed explanations of each scenario.

MSL10 likely functions as a homo-oligomer similar to *Ec*MscS, a homoheptameric channel (Bass et al., 2002; Sukharev, 2002). Thus, the interactions we observe between the N- and the C-terminus could be between domains on the same monomer, or between domains on different monomers once they are assembled into a channel. To distinguish between these two possibilities will require further study. These interactions occur in the absence of the membrane spanning domains, at least in the transient expression system used for BiFC (**Figure 5**). Neither N-terminal phosphorylation status nor the C-terminal lesion S640L had any impact on the self-association of MSL10 in either BiFC or mbSUS assays, suggesting that this is a stable interaction and is not a regulated step in the activation of MSL10 (**Figure 4, 5**).

Based on these results, we propose a three-step model for MSL10 activation, illustrated in **Figure 6**. According to this model, phosphorylation of its N-terminal domain maintains MSL10 in its inactive form (**Figure 6A**). When membrane tension is increased, the channel opens, first resulting in a structural rearrangement at the C-terminus. Second, this conformational change is transduced to the N-terminal domain. Third, the subsequent conformational change of the N-terminus results in its dephosphorylation. The dephosphorylated N-terminal domain (depicted in red) is then capable of triggering a cell death signaling cascade through an unknown mechanism. Dephosphorylation could arise through conformational changes that directly (or indirectly through accessory proteins like 14-3-3 proteins (Moeller et al., 2016)) make the N-terminus more accessible to cytosolic phosphatases or less accessible to kinases. Conformational changes that lead to altered phosphorylation status has been previously shown to activate hormone-receptor complexes in plants (Zhang et al., 2014; Farrell and Breeze, 2018; Wang et al., 2019).

In the case of *MSL10-GFP* overexpression (**Figure 6B**), we propose that the appropriate kinase is not sufficient to maintain the N-terminus in its dephosphorylated form, leading to the accumulation of phosphorylated MSL10-GFP and constitutive activation of the cell death signaling pathway. Phospho-dead versions of MSL10 (MSL10g^7A^) are constitutively active and do not require added tension; conversely, phospho-mimetic version of MSL10 (MSL10g^7D^) are maintained in an inactive state (**Figure 6C**). As for plants harboring the *MSL10^S640L^-GFP* transgene or the *msl10-3G* mutant (**Figure 6D**), the S640L mutation may mimic the conformational change that occurs during opening of the channel, bypassing the normal activation of signaling by membrane tension and leading to constitutive activation. Alternatively, S640L may alter the tension sensitivity or conductivity of MSL10. This effect, however, is blocked by the presence of N-terminal phospho-mimetic substitutions, as shown in plants constitutively expressing *MSL10^7D,S640L^-GFP*. Thus, this three-step model is sufficient to explain the phenotypic and genetic interaction data documented above.

In summary, our three-step model for MSL10 activation, which involves an intra- and/or intermolecular interaction between the soluble N- and C- termini of MSL10, explains the phenotypic similarity shared by plants overexpressing MSL10-GFP, expressing the *MSL10^7A^* transgene, or harboring the *msl10-3G* allele. These results begin to build the groundwork for future investigations into the molecular mechanism by which the mechanosensitive ion channel MSL10 functions.

## ABBREVIATIONS

MS: mechanosensitive;
MSL: MscS-Like;
ROS: reactive oxygen species,
GFP: green fluorescent protein

## SUPPLEMENTARY DATA

**Supplemental Figure 1.** *msl10-3G* is a recessive gain-of-function allele.

**Supplemental Figure 2.** *msl10-3G* allele and *MSL10g^7A^* transgene exhibit similar reduction in fresh weight, petiole length, plant height and apical dominance to *35S:MSL10-GFP.*

**Supplemental Figure 3**. DEX-inducible overexpression of *MSL10* and *MSL10^4A^* promotes the upregulation of MSL10-associated marker genes and ectopic cell death.

**Supplemental Figure 4**. Accumulation of ROS associated with expression of *35S:MSL10-GFP, MSL10g^7A^* transgene and *msl10-3G* allele.

**Supplemental Figure 5**. Incidence of ectopic cell death associated with expression of *35S:MSL10-GFP, MSL10g^7A^* transgene and *msl10-3G* allele.

**Supplemental Figure 6**. Additional controls for the BiFC assay.

**Supplemental Table 1**. List of primers used in this study.

## AUTHOR CONTRIBUTIONS

D.B. and E.S.H. conceived the project and designed the experiments; J.M.S. contributed data to Figures 1C, 3C, and 4; E.S.H. contributed data to Figure 1B and conceived the model in Figure 6. D.B. performed the rest of the experiments. D.B., and E.S.H. wrote the manuscript with input from J.M.S.

## ACKNOWLEDGMENTS

We thank J.-M. Zhou (ShanghaiTech University, Shanghai) for providing *rea1* mutant seeds and the Arabidopsis Biological Resource Center for seeds and for vectors used in BiFC and mbSUS assays. We also are grateful to Ivan Radin, Yanbing Wang, Kari Miller and Angela Schlegel for critical reading of the manuscript. This work was supported by NSF MCB-1253103 and the NSF Center for Engineering Mechanobiology Award 1548571. J.M.S. was supported by NSF Graduate Research Fellowship DGE-1745038. No conflicts of interest declared.

---

**Supplemental Figure 1.**
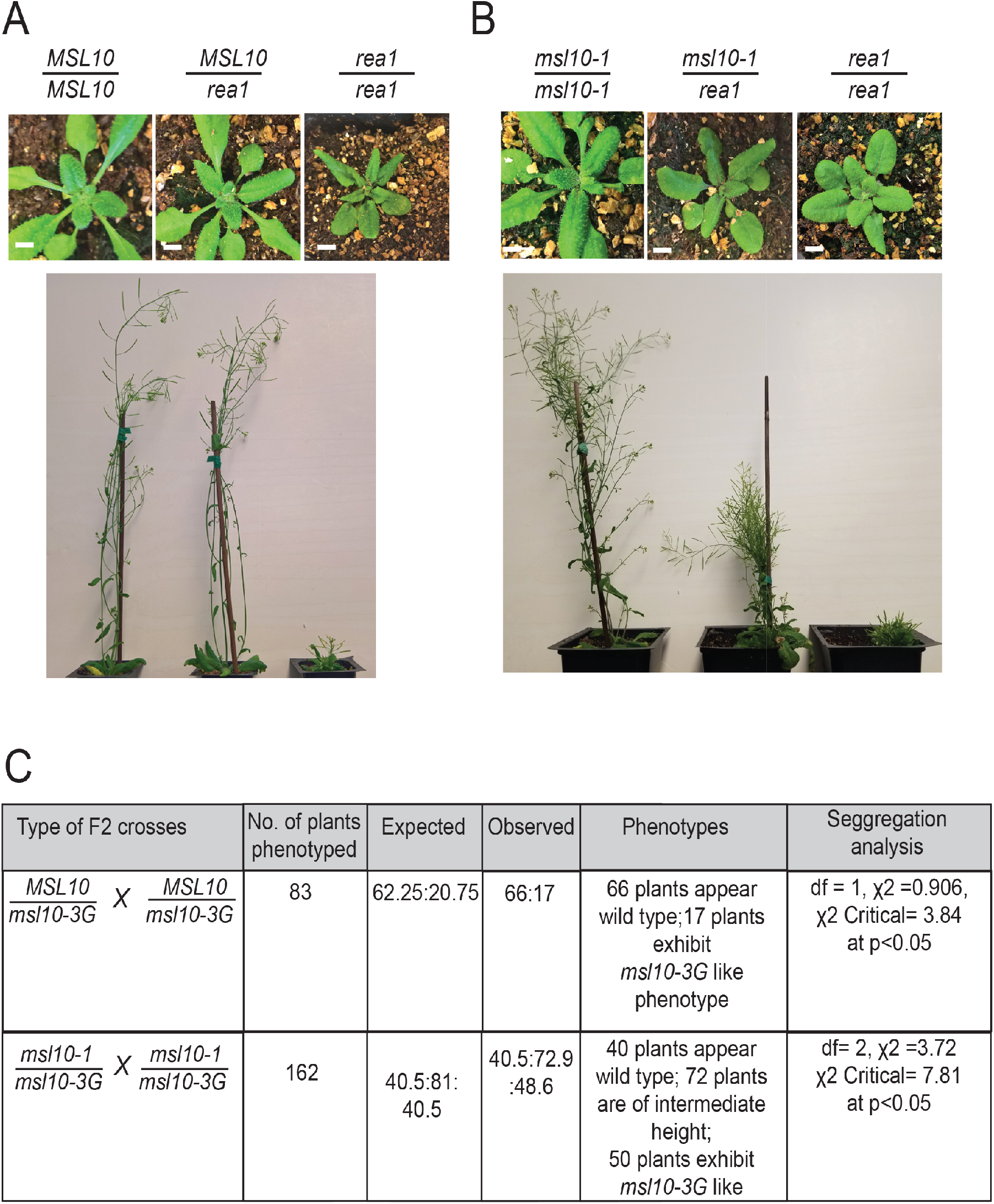
*msl10-3G* is a recessive gain-of-function allele. (A) Representative images of parental and F1 offspring plants from a *msl10-3G* backcross to wild type *Col-0.* Top, four-week-old; bottom, eight-week-old plants. Bar = 0.5 cm. (B) Representative images of parental and F1 offspring from a *msl10-3G* outcross to *msl10-1.* Top, four-week-old plants; bottom, twelve-week-old plants. Bar = 0.5 cm. (C) Table showing segregation of *msl10-3G* inheritance data compared to the expected ratios using a chi-squared analysis. Plants were randomly chosen for genotyping.

**Supplemental Figure 2.**
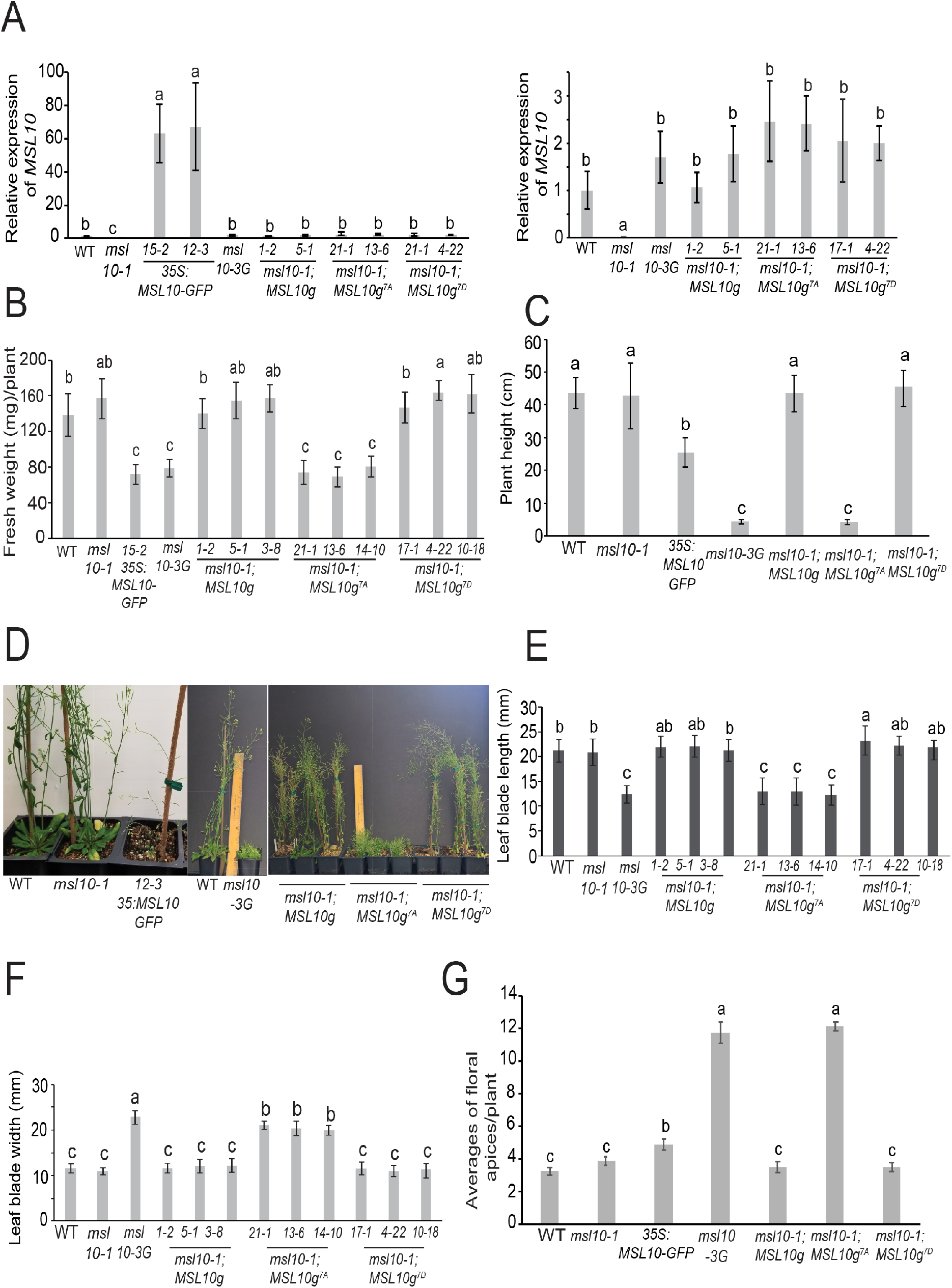
*msl10-3G* allele and *MSL10g^7A^* transgene exhibit similar reduction in fresh weight, petiole length, plant height and apical dominance to *35S:MSL10-GFP*. **(A)** Relative transcript abundance of *MSL10* in the indicated lines. Left panel displays accumulation of *MSL10* transcripts in plants harboring the *35S:MSL10-GFP* transgene, genomic *MSL10g* phospho-variants, the EMS-induced point mutant allele *msl10-3G* or the null mutant allele *msl10-1* compared to wild-type *Col-0*. Right panel depicts *MSL10* expression only in wild type, null mutant allele and genomic *MSL10g* phospho-variants. cDNA was synthesized from total RNA extracted from rosette leaves of three-week-old plants. Expression levels of respective genes were analyzed with qRT-PCR and normalized to both *EF1a* and *UBQ5.* Mean fold-change values relative to the wild type are plotted, with error bars showing ±SE of the mean of three biological replicates. **(B)** Average fresh weight of rosettes clipped from three-week-old plants of the indicated genotypes. Error bars indicate ±SD of three replicates (n=10 per replicate). **(C)** Plant height measurement from eight-week-old plants from indicated genotypes. Error bars indicate ±SD of three replicates (n=20-30 per replicate). Different letters indicate significant difference as determined by one-way ANOVA followed by Scheffe’s post-hoc test (P < 0.05) for unbalanced samples. **(D)** Representative images of eight-week-old adult plants from indicated genotypes. **(E)** Leaf blade length of seventh leaf from 35-day-old plants from the indicated genotypes. Error bars indicate ±SD of three replicates (n=18 per replicate). **(F)** Leaf blade width of fourth or fifth leaf from 35-day-old plants from the indicated genotypes. Error bars indicate ±SD of three replicates (n=18 per replicate). **(G)** Average number of floral apices were counted from six to seven-week-old plants of the indicated genotypes. Error bars indicate ±SD of three replicates (n=20 per replicate). The T4 homozygous transgenic lines expressing either *35S:MSL10-GFP* in Col-0 background (*15-2)* and T3 homozygous transgenic lines expressing *MSL10g,* or *MSL10g^7D^* in the *msl10-1* background, and T3 segregating or T3 homozygous lines for *MSL10g^7A^* were selected for comparison. Plants of indicated genotypes were grown side-by-side in soil at 21°C under a 24-h light regime. Different letters indicate significant differences, as determined by one-way ANOVA followed by Tukey’s post-hoc test (P < 0.05).

**Supplemental Figure 3.**
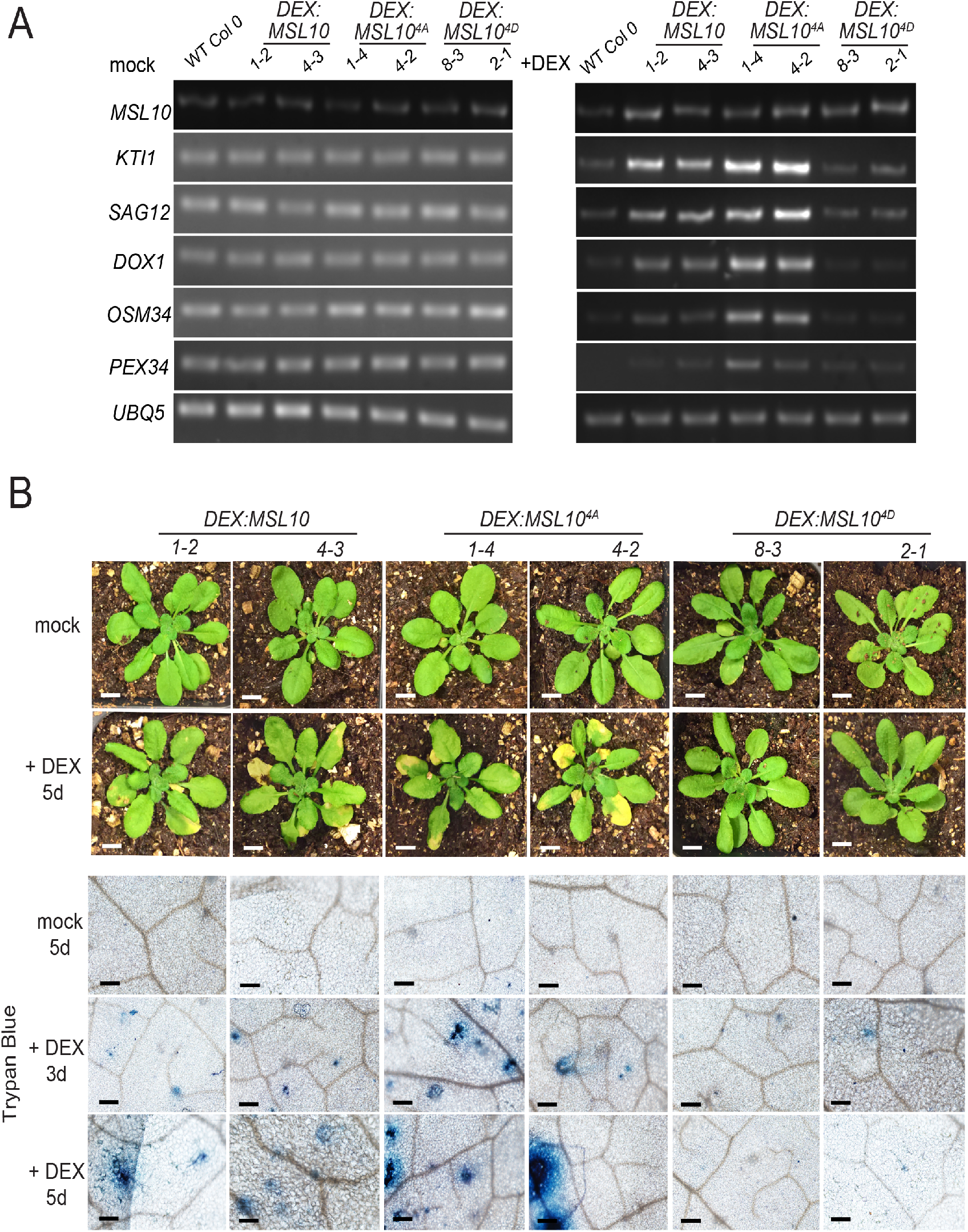
DEX-inducible overexpression of *MSL10* and *MSL10^4A^* promotes the upregulation of MSL10-associated marker genes and ectopic cell death. Representative images of five-week old plants grown under a short-day photoperiod (10 h light/14 h dark) sprayed with 0.016% ethanol (mock; top row) or 30 μM DEX (bottom row) for 5 days. Expression profiles of five marker genes before and after DEX treatment of transgenic plants expressing *DEX:MSL10* and its phospho-variants using semi-quantitative RT-PCR. Rosette leaves from five-week-old plants were infiltrated with 10 μM DEX or mock (0.016% ethanol), and tissue was harvested 12 h post infiltration. Total RNA was isolated from *Col-0*, and from two independent homozygous T3 lines expressing either *DEX:MSL10, DEX:MSL10^4A^* or *DEX:MSL10*^4D^. Expression of *UBQ5* was used as a control.

**Supplemental Figure 4.**
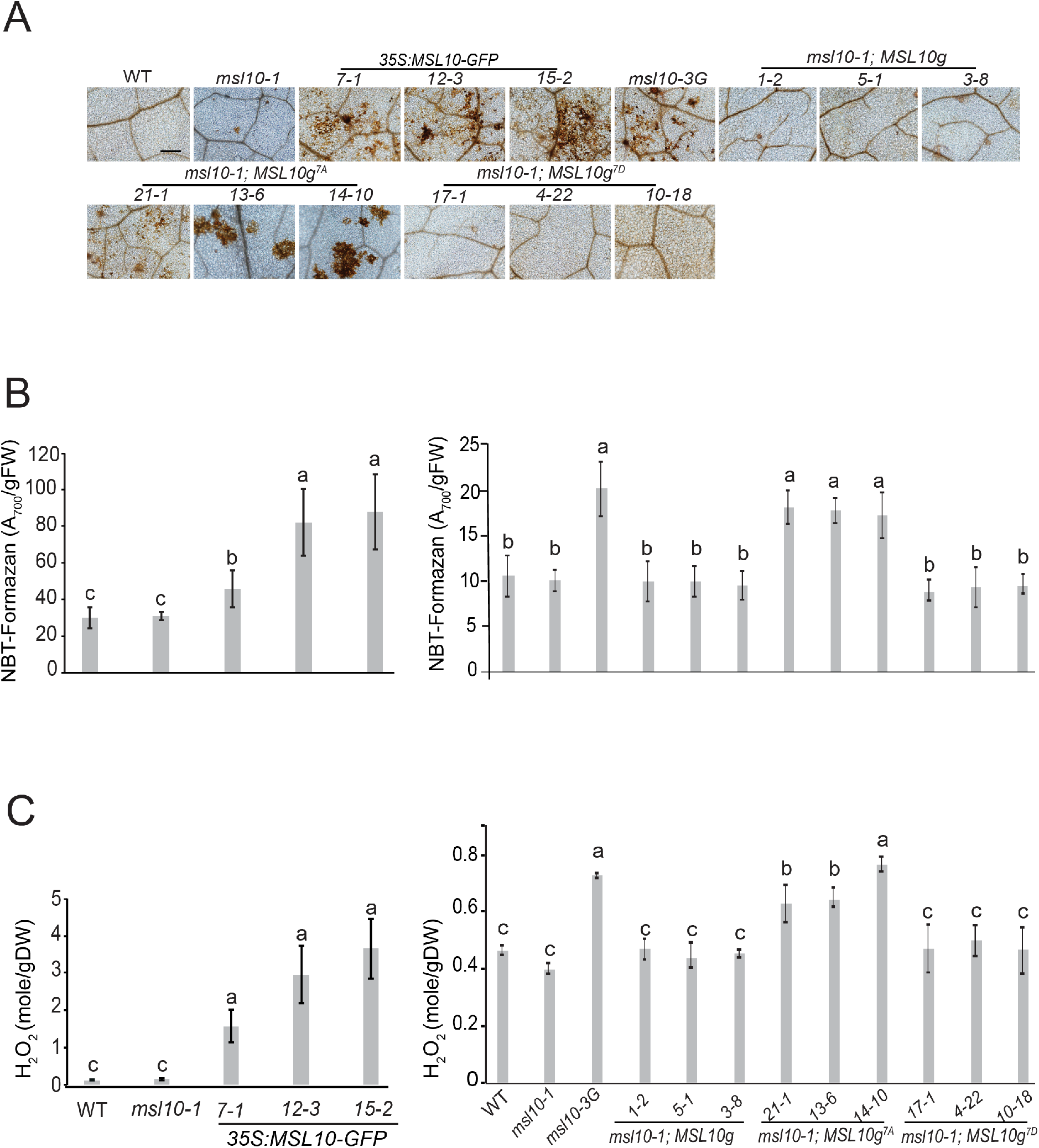
Accumulation of ROS associated with expression of *35S:MSL10-GFP, MSL10g^7A^* transgene and *msl10-3G* allele. **(A)** Elevated levels of H_2_O_2_ detected by DAB staining. Representative bright-field images of rosette leaves are shown. Bar = 200 μm. **(B)** Colorimetric quantification of NBT-formazan deposition on rosette leaves. FW, fresh weight. Data are means ± SD of three replicates (n=18). **(C)** Hydrogen peroxide (H_2_O_2_) content, quantified as the H_2_O_2_ (mmol) per gram dry weight (DW) of rosette leaves using Amplex Red-coupled fluorescence quantitative assay. Values are means ± SD of three replicates (n=14). **(A), (B)**, and **(C)** Leaves from three-week-old plants from each line grown at 21°C (top row) under 24-h light regime were used for the experiment. Three independent T3 homozygous transgenic lines expressing *MSL10g, MSL10g^7A^* or *MSL10g^7D^* in the *msl10-1* background were selected for comparison.

**Supplemental Figure 5.**
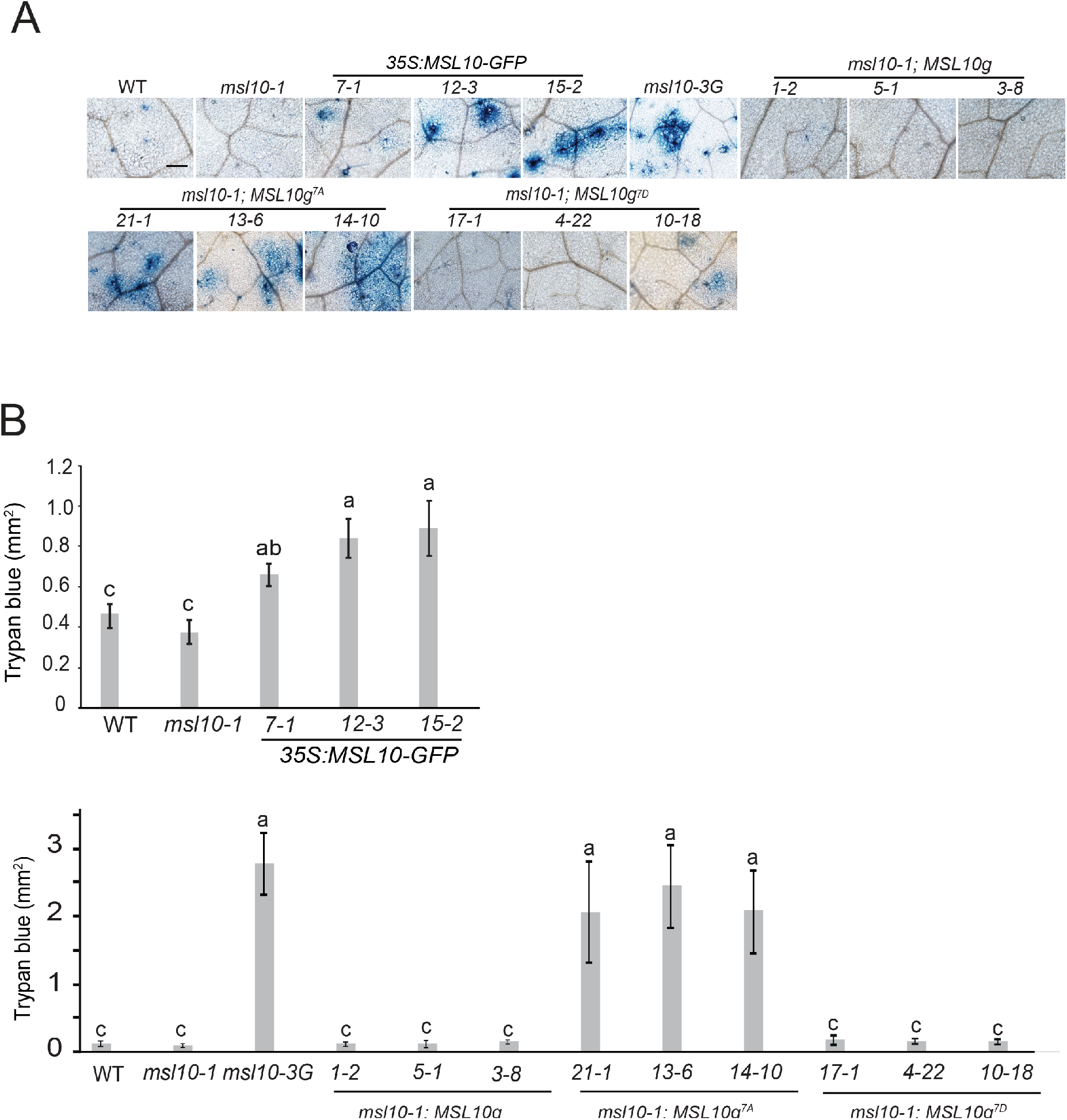
Incidence of ectopic cell death associated with expression of *35S:MSL10-GFP, MSL10g^7A^* transgene and *msl10-3G* allele. **(A)** Visualization of ectopic cell death incidence by Trypan blue staining. Bar = 200 μm. **(B)** Quantification of ectopic cell death detected by Trypan blue staining. Data are means ± SD of three replicates each one consisting of 7 to 10 leaves. **(A)** and **(B)** Leaves from three-week-old plants from each line grown at 21°C (top row) under 24-h light regime were used for the experiment. Three independent T3 homozygous transgenic lines expressing *MSL10g, MSL10g^7A^* or *MSL10g^7D^* in the *msl10-1* background were selected for comparison.

**Supplemental Figure 6.**
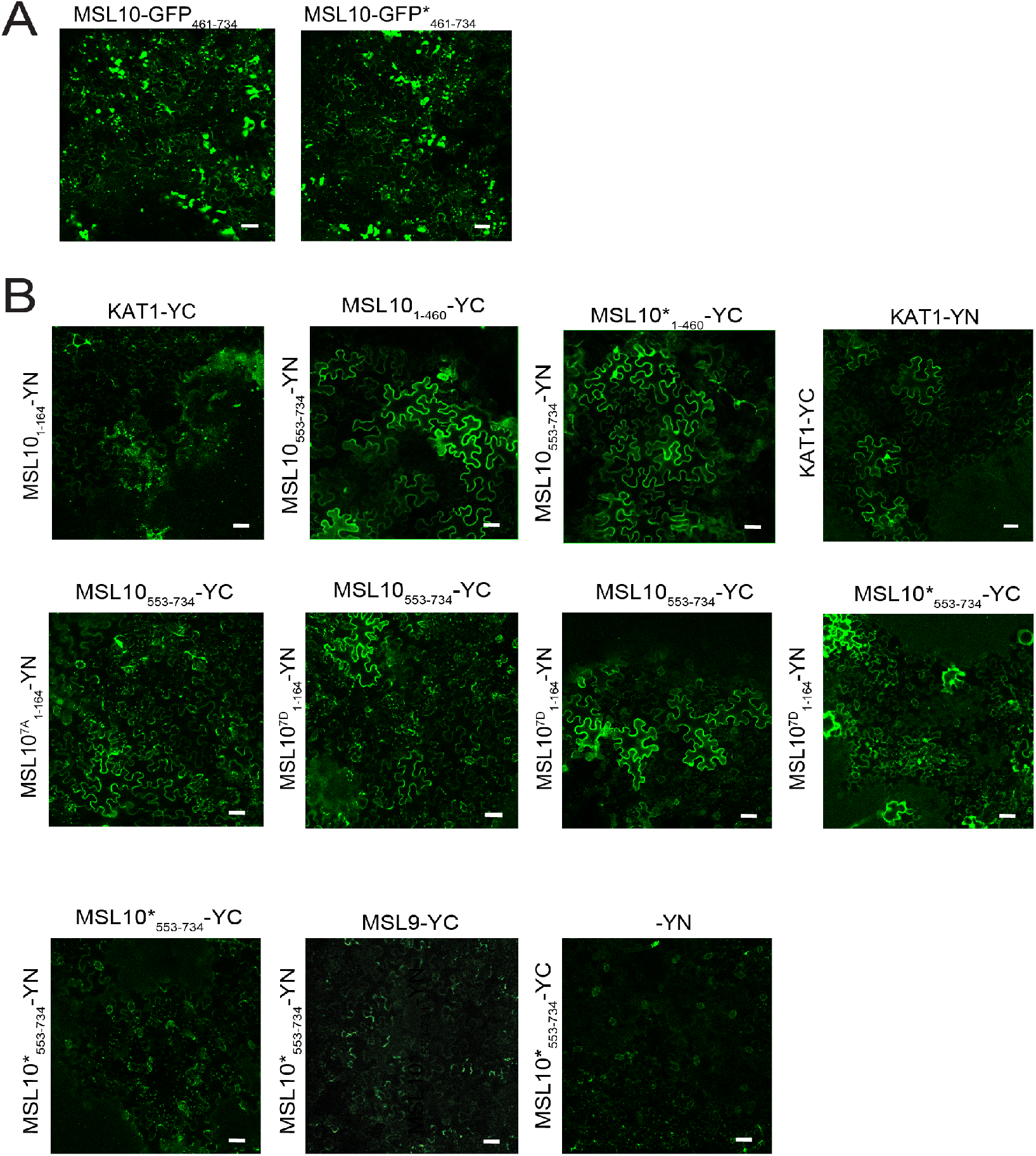
Additional controls for the BiFC assay. **(A)** *N. benthamiana* leaves co-infiltrated with constructs fused to the split YFP were observed under the confocal microscope 3-5 days after infiltration. Scale bar, 50 μm. (Top row) Neither domain interacts with KAT1. (Middle row) Neither phosphorylation status of the N-terminus nor the S640L lesion in the C-terminus affect the interaction. either reconstituted YFP fluorescence signal from positive interaction or marked reduction in YFP signal in case of negative interaction from indicated combinations. (Bottom row) Negative controls for the N-terminus with the S640L lesion. Scale bar, 50 μm. The C-terminal half of MSL10 forms aggregates when fused to GFP. Representative confocal images of abaxial *N. benthamiana* leaf epidermis from plants transiently expressing truncated MSL10_461-734_-GFP and MSL10 ^S640L^_461-734_-GFP. Images were taken 5 d postinfiltration. Scale bar, 50 μm.Supplemental Table 1. List of primers used in this study.

**Supplemental Table 1.**
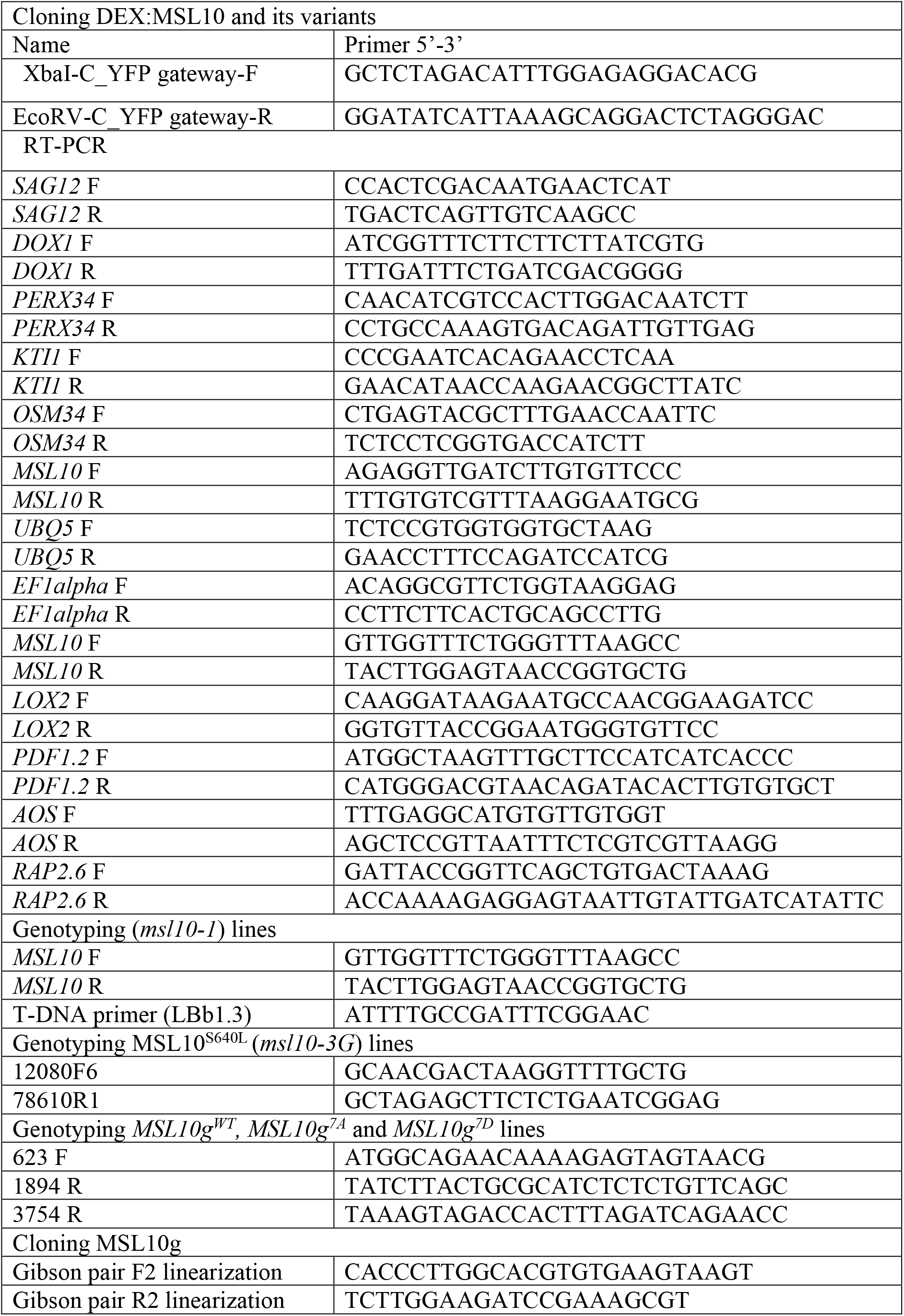
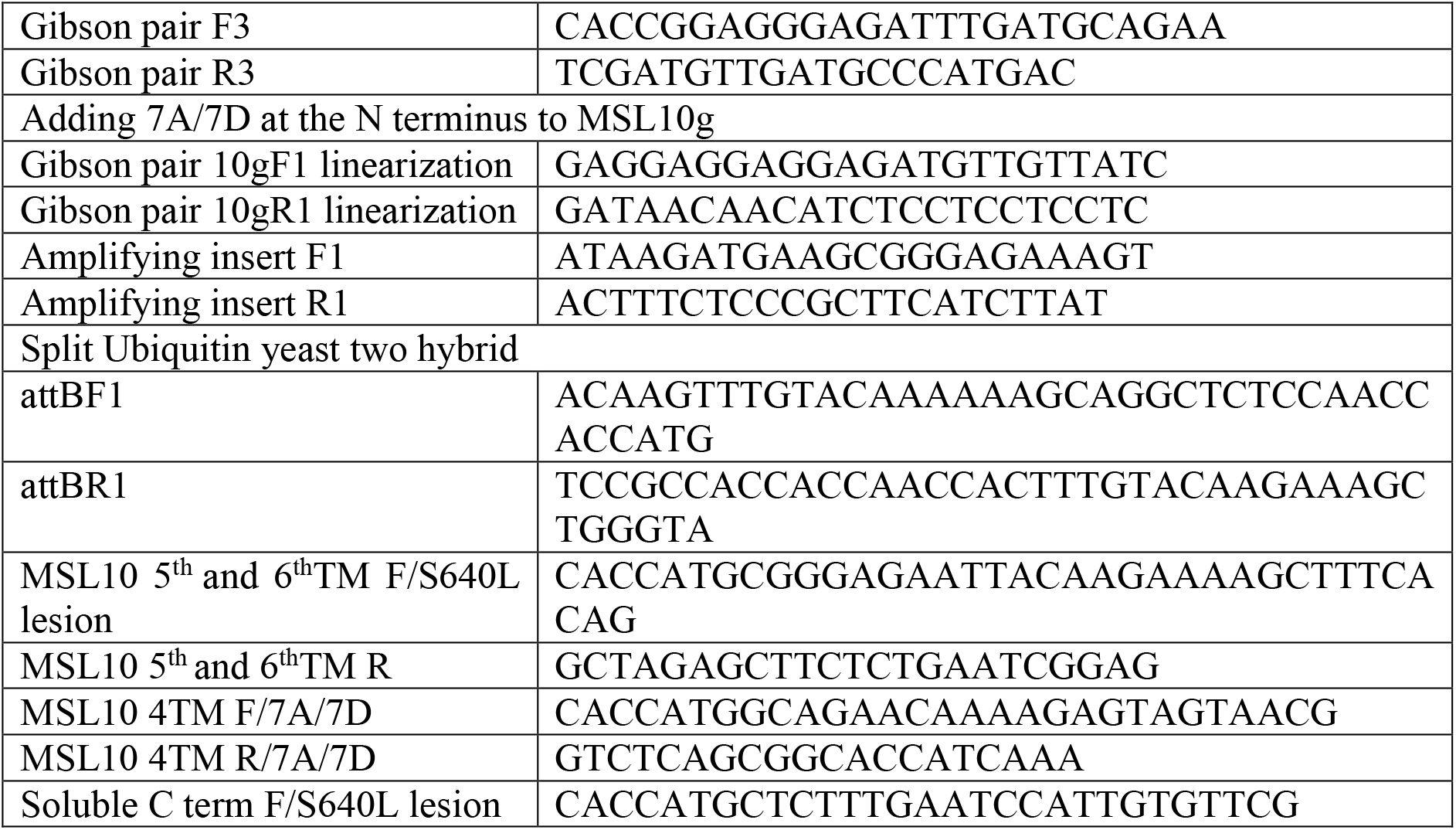
List of primers used in this study.

